# Task Splitting and Prompt Engineering for Cypher Query Generation in Domain-Specific Knowledge Graphs

**DOI:** 10.1101/2025.04.23.649790

**Authors:** Saber Soleymani, Nathan Gravel, Krzysztof Kochut, Natarajan Kannan

## Abstract

The integration of large language models (LLMs) with knowledge graphs (KGs) holds significant potential for simplifying the process of querying graph databases, especially for non-technical users. KGs provide a structured representation of domain-specific data, enabling rich and precise information retrieval. However, the complexity of graph query languages, such as Cypher, presents a barrier to their effective use by non-experts. This research addresses the challenge by proposing a novel approach, Prompt2Cypher (P2C), which leverages task splitting and prompt engineering to decompose user queries into manageable subtasks, enhancing LLMs’ ability to generate accurate Cypher queries that align with the underlying graph database schema. We demonstrate the effectiveness of P2C in two biological KGs (protein kinase and ion-channel) that differ in size, schema and complexity. Compared to a baseline approach, our method improves query accuracy, as demonstrated by higher Precision, Recall, F1-score, and Jaccard similarity metrics. This work contributes to the ongoing efforts to bridge the gap between domain-specific knowledge graphs and user-friendly graph database query interfaces.

## Introduction

Knowledge graphs (KGs) are effective data representation systems that allow integration and conceptualization of information from diverse sources and formats^1^. KGs provide a rich framework for knowledge discovery and data mining by representing information as entities (nodes), relationships (edges), and attributes. In recent years, domain-specific KGs, such as those in biology or chemistry, have become increasingly important for advancing research in these fields by structuring and connecting vast amounts of specialized information^2^. This study examines two KG examples which are shown in Figure 1: a. ProKinO (Protein Kinase Ontology)^3^, a well-established KG in the field of protein kinases, which stores, integrates, and represents diverse protein kinase-related data and their relationships, and b. The Ion Channels Knowledge Graph (ICKG), which captures data related to ion channel proteins introduced in this study.

**Figure 1.**
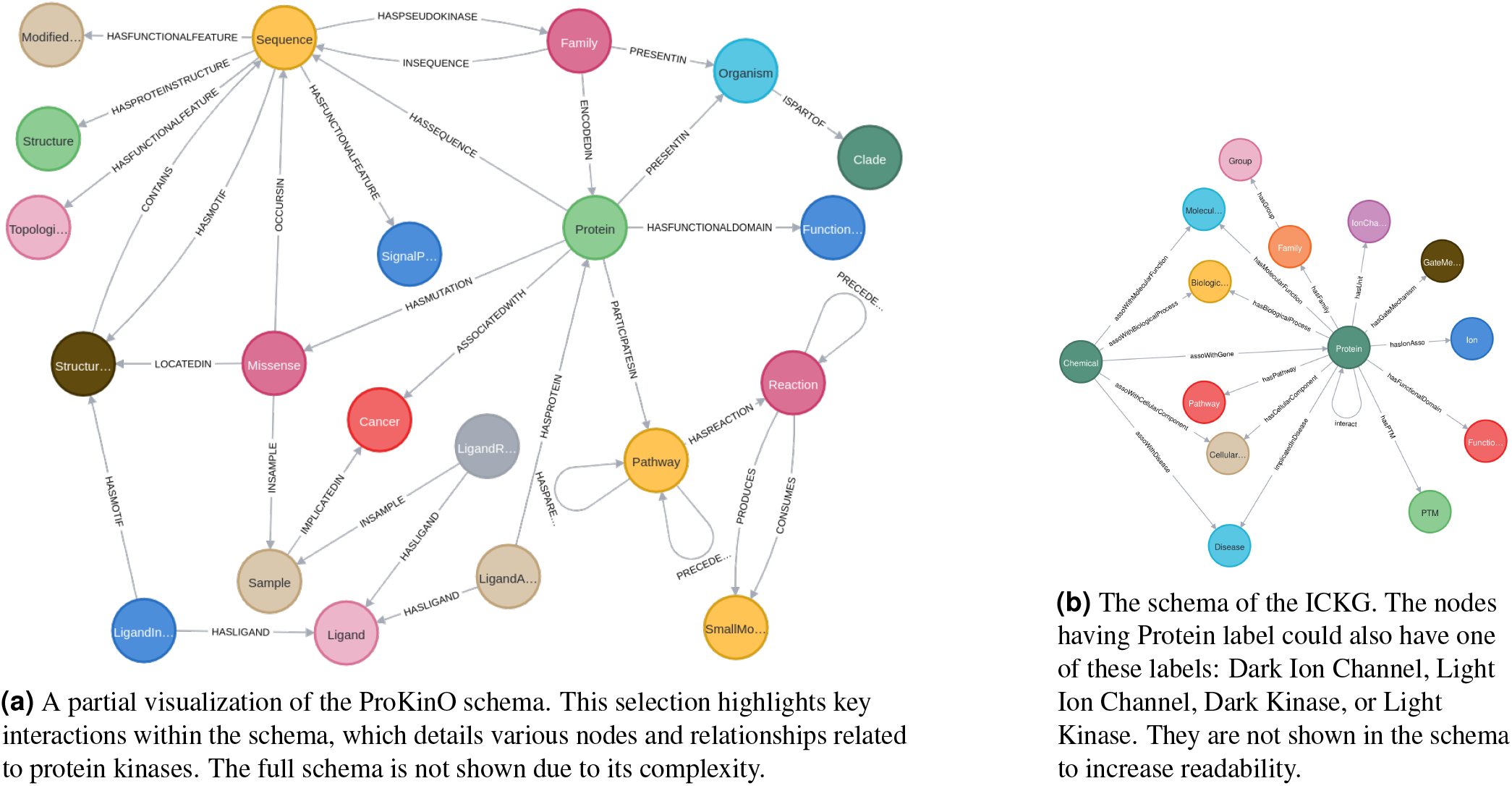
Comparison of the ICKG and ProKinO knowledge graph schemas.

In a practical application, such as studying ion channels, a KG allows researchers to explore the intricate relationships between various biological entities^4^. For example, they can identify which proteins are involved in specific pathways, how different ion channels interact with each other, or how certain chemicals are linked to diseases. This structured representation of knowledge facilitates advanced queries, data integration, and hypothesis generation. Despite their potential, the complexity of these graphs often presents significant challenges for users who are not experts in graph query languages. Specialized query languages such as SQL for relational databases, allow users to navigate the interconnected structures of data. While these languages enable precise pattern matching, their syntax complexity creates barriers for users^5^. Therefore, the integration of natural language interfaces (NLIs) with these databases, also known as Natural language interfaces to databases (NLIDBs), is a rational way to make data navigation more accessible^5–7^. Similarly, query languages such as SPARQL or Cypher are used to extract information from graphs. Users, especially domain experts in fields such as biology, face the same challenge of crafting complex patterns of nodes and relationships to retrieve relevant data. Crafting such queries is challenging for non-technical users due to the intricacies of the syntax and the need to understand the underlying graph structure^8^.

NLIs and NLIDBs have been studied for decades, initially relying on rule-based or grammar-driven methods^9^. Early systems defined hand-crafted grammars or logic-based formalisms to map natural language to queries. For example,^10^ employed combinatory categorial grammar (CCG) to parse sentences into logical forms. A major focus within NLIDBs has been text-to-SQL research, which seeks to translate natural language questions into SQL queries^11^. These classical methods often required extensive domain tuning. Subsequent efforts incorporated database semantics via ontologies or query logs to improve interpretation^9^, or used methods such as intermediate user feedback to refine queries^9,12^. These early approaches demonstrated the feasibility of natural language querying, but were limited by pre-defined rules and the need for manual engineering for each new domain or schema.

LLMs have demonstrated impressive capabilities in understanding and generating human language^13^, as well as code^14^, protein sequences^15^, and other textual data^13^. However, LLMs often hallucinate when dealing with domain-specific knowledge not covered extensively in their training data^16,17^. This problem arises because LLMs generate responses based on learned patterns, which may not always align with specific domain knowledge^18^. This limitation is particularly pronounced in fields like biology and health care^19^, which rely heavily on structured formats such as knowledge graphs. Consequently, integrating LLMs with this structured domain knowledge holds significant promise for advancing research and technological applications. Despite their advancements, LLMs struggle with domain-specific queries, particularly those involving complex knowledge graphs^20^. More recently, deep learning approaches have transformed NLIDB by treating query translation as a machine translation problem, converting natural language into formal queries using encoder-decoder models^9,21–23^. This shift towards data-driven approaches was motivated by the ability of sequence-to-sequence models to learn complex mappings directly from large datasets without requiring explicit rule engineering. However, their auto-regressive decoding can struggle with very complex or long queries, often making mistakes on multi-hop query logic or when composing many conditions^9^. These limitations motivated explorations into alternative generation paradigms and the use of larger pre-trained language models.

Recent advances in deep learning have shifted the focus toward applying LLMs to query translation tasks, but these models face unique challenges when generating structured queries. In the context of Text-to-Cypher tasks, hallucinations manifest in several specific ways that directly impact query validity and execution. We define hallucinations in this domain as instances where the LLM generates: (1) Non-existent node labels (e.g., generating “GeneSequence” when no such node type exists in the schema); (2) Invalid relationship types (e.g., using “INTERACTS_WITH” when the correct relationship is “hasInteraction”); (3) Incorrect property names (e.g., referencing “name” property when the schema uses “hasPrimaryName”); (4) Schema structure misalignment (e.g., attempting to connect nodes that have no direct relationship in the schema); or (5) Syntactically valid but semantically incorrect queries that fail to address the user’s question.

These hallucinations arise when LLMs attempt to generate Cypher queries without fully understanding or properly navigating the knowledge graph’s schema, particularly in domain-specific contexts where terminology and relationships are highly specialized. This difficulty occurs because LLMs, primarily trained on unstructured text, struggle to grasp the inherent structure and relationships within a knowledge graph. They may misinterpret relationships between entities, fail to identify the correct properties to query, or generate syntactically incorrect queries that do not align with the KG’s schema. This challenge highlights the need for more accessible and accurate tools that bridge the gap between complex data structures and user queries, enabling easier interaction with domain-specific knowledge through LLMs.

LLMs, due to their large-scale pre-training and unique ability to understand texts, introduce approaches that can understand the schema and user’s needs better. The application of LLMs to Text-to-SQL represents perform better compared to earlier approaches, as these models can leverage their pre-trained knowledge to understand and generate more complex and nuanced SQL queries^11^. Building on these advances in Text-to-SQL, researchers have begun exploring similar approaches for Text-to-Cypher translation. Datasets, such as Spcql^24^ collect pairing natural language with Cypher queries, provide essential resources for this research direction. Techniques such as zero-shot and few-shot prompting are being investigated to leverage the pre-trained knowledge of LLMs to translate natural language queries into Cypher without requiring extensive task-specific training data^25^. Additionally, fine-tuning LLMs on datasets of natural language queries paired with their corresponding Cypher queries is being explored as a way to further improve the performance of this translation task^25^. The effectiveness of fine-tuning often depends on the availability of high-quality, task-specific datasets, which remains an area of ongoing development in the Text-to-Cypher domain^25^.

Within the broader context of graph databases and knowledge representation, the ProKinO knowledge graph^3^ exemplifies the conceptualization of complex relationships among protein kinase sequence, structure, function, and disease in a format that is both human-readable and machine-readable. While Text2Cypher^25^ is effective for simpler graph schemas such as movie databases, we found that such general-purpose approaches struggle with complex domain-specific knowledge graphs due to schema heterogeneity, diverse node and relationship types, and multi-hop queries. When evaluated on ProKinO and ICKG, Text2Cypher failed to answer any of our test questions correctly (0% accuracy), emphasizing the need for domain-adapted solutions. Recent analyses have shown that even fine-tuned Text2Cypher models exhibit significant performance degradation when applied to complex schemas in less common domains with limited training data^26^.

This is also shown in other research such as^27^, which demonstrated that LLMs exhibit different levels of prior knowledge across query languages, with performance correlating to the prevalence of these languages in training data. Their work showed that while LLMs performed reasonably well on SQL with minimal schema information, other languages like SPARQL required more explicit schema context and examples.

LLMs demonstrate enhanced performance and reduced hallucination when provided with relevant contextual information during generation. Techniques like Retrieval-Augmented Generation (RAG) aim to supply this context by retrieving relevant unstructured text passages from external knowledge sources, thereby grounding the model’s output in factual data, especially for knowledge-intensive tasks where parametric knowledge alone may be insufficient or outdated. However, for tasks involving structured data like knowledge graphs, the primary context is often the knowledge graph’s schema itself, defining the valid nodes, relationships, and properties. Yet, domain-specific KGs, particularly in fields like bioinformatics as explored in this work, can possess large and intricate schemas (e.g., ProKinO with dozens of node/edge types). Presenting the entire schema to an LLM can be counterproductive due to context window limitations and the risk of overwhelming the model with irrelevant information, potentially hindering its ability to identify the correct entities and paths for a given query.

In Text-to-Cypher research, SyntheT2C^28^ proposed a methodology for constructing synthetic Question-Cypher pair datasets to fine-tune LLMs on the Text-to-Cypher task. Their approach combines two complementary pipelines: LLM-based prompting using GPT-4o and template-filling to generate diverse syntactic patterns. While this approach successfully creates large-scale training data (their MedT2C dataset contains 3,000 samples) without requiring manual annotation, our work takes a different approach by focusing on a curated set of domain-specific test queries that have been verified by subject matter experts to represent realistic scenarios that biologists might encounter when querying specialized knowledge graphs like ProKinO and ICKG.

Several approaches have been proposed to address the challenge of translating natural language into accurate and executable graph queries. Template-based systems^29^ and fine-tuning^30^ have been used to map natural language queries to SPARQL or Cypher. Other works explore the use of LLMs for tasks such as knowledge graph completion^31^, link prediction^32^, relation classification^33^, and reasoning^34^. However, these methods often require extensive training data, computational resources, or are limited to specific tasks within the KG domain. Notably, these methods do not focus on generating executable graph queries like Cypher from natural language inputs. They also do not accommodate the complexities of domain-specific KGs.

The limitations of general-purpose NLIs underscores the need for specialized approaches that can handle the diverse relationships and domain-specific constraints inherent in biomedical knowledge graphs.

Our approach differs from existing methods by leveraging LLMs to generate accurate Cypher queries from natural language inputs without requiring additional training or fine-tuning. However, LLMs are inherently designed to process sequential data^35^, making it challenging to navigate the complex structures of KG schemas, including node labels, properties, and relationships. We address these challenges by employing two key strategies: task splitting and prompt engineering^36^, which enable LLMs to effectively generate queries aligned with domain-specific KGs and reduce hallucinations.

Our research explores two key aspects to customize LLMs for querying structured data like KGs: task splitting and prompt engineering. Task splitting involves breaking down a complex task into smaller, manageable subtasks. This allows the LLM to focus on understanding specific components of the query, such as identifying relevant entities, relationships, and attributes within the KG’s schema. Our method splits query generation into node identification and Cypher synthesis. This strategy narrows the LLM’s focus to schema-relevant components, improving precision in multi-hop queries.

We demonstrate the effectiveness of our method using two domain-specific knowledge graphs, ProKinO and ICKG, as examples. We specifically chose these KGs because they represent different levels of complexity and domain-specific focus. ProKinO, a well-established KG in the field of protein kinases, provides a robust testbed with diverse entities and relationships. ICKG, a novel KG focusing on ion channels introduced in this research, offers a contrasting case with a more focused domain and specific query requirements. Our experiments show that our approach improves query accuracy compared to baseline methods, which involve directly asking an LLM to generate a Cypher query based on a user query, the KG’s schema, and general instructions. Specifically, our method reduces the occurrence of hallucinations and generates queries that are more aligned with the KG’s structure. To evaluate the effectiveness of our approach, we use precision, recall, F1-score, and Jaccard similarity metrics to compare the set of results of ground truth Cypher queries with the results of executing generated Cypher queries.

## Methods

### Task Description

The knowledge graph *G* is a labeled property graph (LPG), loaded into a Neo4j graph database, represented as a directed graph *G* = (*V, E*), where *V* is the set of nodes (entities) and *E* is the set of edges (relationships). Our primary task is to convert a user’s natural query *Q* into an executable Cypher query *C* that retrieves information from *G*. The process involves first identifying a subset of relevant nodes *N*^′^ ⊆ *V* and their associated relationships *E*^′^ ⊆ *E* before generating the final Cypher query *C*. This approach allows us to manage the complexity of large knowledge graphs by considering only the most relevant portions of the schema for a given query, while leveraging schema comments to enrich the semantic understanding of the graph structure.

### Preprocessing and KG generation

We used two knowledge graphs (KGs) as case studies to generate Cypher queries from natural language inputs: Protein Kinase Ontology (ProKinO)^3^ and Ion Channel Knowledge Graph (ICKG). Both KGs were loaded into a Neo4j graph database, enabling us to run Cypher queries for data retrieval and analysis.

ProKinO is a knowledge graph designed to conceptualize and unify diverse protein kinase data related to sequence, structure, function, and disease. Originally developed to support integrative analysis and hypothesis generation in kinase-focused research, ProKinO has since been expanded to incorporate additional classes, relationships, and datasets, including kinase expression patterns and drug interaction information^3^. Comprising 7,430,379 nodes of 47 distinct types, such as Protein, GeneExpression, Ligand, and Pathway, and 19,110,051 edges of 37 types, including relationships such as proteins to functional domains, pathways to reactions, and ligand interactions to structural motifs, ProKinO is a comprehensive resource for exploring well-studied (light) and understudied (dark) kinases^37^. The knowledge graph supports complex SPARQL queries for various tasks such as linking kinases to associated pathways and disease phenotypes. This rich semantic context makes ProKinO particularly suitable as a test bed for evaluating natural language to Cypher query generation. The extensive interconnectedness of entities within this KG poses a challenge for LLMs, which may struggle to identify the most relevant paths and properties for a given query.

We developed the ion channels knowledge graph (ICKG) by integrating data from repositories such as UniProt^38^, STRING^39^, The Gene Ontology^40^, and Reactome^41^. It comprises 1,064,666 nodes of 18 distinct types, such as Protein, Pathway, and Functional Domain, and 6,288,620 edges of 18 types, including relationships such as proteins (including light and dark ion channels) to biological processes, diseases, ions, and PTMs. To construct this knowledge graph, we reviewed and incorporated 429 curated ion channel protein identifiers from UniProt. We further integrated protein-protein interaction (PPI) data from the STRING database^39^, filtering for high-confidence scores (≥ 700) to add 5,062 new PPI edges related to ion channels. Additionally, we added 12,542 new edges related to gene ontology relationships to all additional ion channel nodes from The Gene Ontology database^40,42^. Also, 296 reaction relationships were added to existing lowest level Reactome pathway nodes retrieved from the Reactome database^41^. Other relationships associated with edge types such as *hasGroup, hasFamily, hasUnit, hasGateMechanism*, and *hasIonAsso* are obtained from a manual curation gathered from a ion channel specific classification^4^. We developed a Python-based ETL (Extract, Transform, Load) pipeline. This pipeline was used to batch-process nodes and edges into a Neo4j database. By assigning appropriate labels to entities and forming relationships based on domain standards, we improved the utility of ICKG for complex queries and data analysis in ion channel research.

### Proposed Approach: Task Splitting and Prompt Engineering

To overcome the limitations of direct query generation, we developed a two-pronged approach that leverages task splitting and prompt engineering to enhance the accuracy of LLM-generated Cypher queries. This approach systematically breaks down the complex process of query generation while providing structured guidance to the LLM at each step.

Research has shown that LLMs perform more effectively when prompted with specific, smaller tasks^36,43^. Our approach decomposes the query generation process into two main subtasks: (1) identifying relevant nodes from the schema based on the user’s query, and (2) generating the final Cypher query using the filtered subgraph and metadata. For the first subtask, we prompt the LLM to analyze the natural language query and determine which entities within the schema are pertinent. The system then automatically filters relationships to include only those involving the identified nodes, creating a focused subgraph that simplifies the query generation process.

The filtered subgraph serves as a constrained reasoning space that mirrors human expert query formulation. By first identifying relevant nodes (e.g., Protein, Organism) and their immediate relationships, the system establishes anchor points for multi-hop reasoning. This filtering process reduces hallucination risks by: 1) eliminating irrelevant schema elements that could distract the LLM, and 2) creating a tractable search space for relationship path discovery. The system identifies optimal paths through this filtered subgraph using a three-step process: (1) using the LLM to identify relevant nodes from the available node labels based on query analysis, (2) filtering relationships to only those involving the identified nodes, and (3) leveraging nodes’ descriptions, properties, and relationship metadata to guide the LLM in generating appropriate Cypher queries. Rather than employing explicit pathfinding algorithms, we leverage the LLM’s semantic understanding to implicitly determine the most appropriate path through the filtered subgraph, ensuring alignment with both the user’s intent and the KG’s actual connectivity patterns.

For each subtask that interacts with the LLM, we design tailored prompts that provide clear instructions and relevant context. The context includes the filtered schema, node descriptions, properties, and relationship metadata. This structured prompting approach ensures that the LLM can focus on specific aspects of the query with relevant context, thereby reducing the likelihood of errors and hallucination when generating Cypher queries. In the first subtask, the prompt is crafted to guide the LLM in identifying the relevant start and end node labels based on the user’s natural language query and the knowledge graph schema. When generating the final Cypher query, the prompt integrates the identified nodes, selected path, and extracted properties, instructing the LLM to construct a syntactically correct and executable Cypher statement that aligns with the knowledge graph’s schema and accurately retrieves the requested information.

To illustrate our prompting approach, Prompt1 follows this template: “Return a list of nodes needed to run this query: ‘{user_query}’. The relevant nodes must be from this list: {list_of_nodes}”. After programmatically extracting relationships and properties for the identified nodes, Prompt2 uses this structure: “Generate a Cypher query to answer: ‘{user_query}’. Consider these relevant nodes ({relevant_nodes}) with relationships: {relevant_relationships}. Each node includes properties such as {node_properties} and descriptions like {node_descriptions}. Follow these constraints: {cypher_instructions}”. These prompts guide the LLM through a focused, two-step reasoning process that reduces hallucinations by constraining the generation space to relevant schema elements.

The development of our approach followed an iterative process to optimize query generation accuracy. The process began with reformulating test queries to ensure explicit mapping to the schema’s navigable pathways. We then implemented task splitting to decompose query generation into manageable subtasks, allowing focused processing of each component. These steps were refined through careful prompt engineering, with instructions crafted to guide the LLM in producing accurate Cypher queries.

The node selection process follows a systematic approach to identify relevant entities from the schema. Rather than using traditional keyword matching or rule-based methods, our approach leverages the natural language understanding capabilities of LLMs to identify semantically relevant nodes. The process begins by extracting all available node labels from the knowledge graph schema, which provides the complete vocabulary of possible entities that can be referenced in queries. This set of labels serves as a constraint to ensure that only valid nodes are selected. We then construct a prompt that combines the user’s natural language query with the extracted node labels. This prompt guides the LLM to analyze the semantic relationship between the query and the set of KG labels available in the schema. For example, when processing a query like “Find all dark ion channels that bind to calcium,” the LLM would identify ‘DarkIonChannel’ and ‘Ion’ as the relevant nodes, creating a focused subset of the schema for query generation.

Our implementation uses a modular approach to ensure flexibility and maintainability. The system first processes the user query to identify relevant nodes, then filters the schema to include only those nodes and their associated relationships. For each node, we include its description, properties, and relationships as context in the LLM prompt. This filtered context is crucial for managing the token limitations of LLMs while providing the most relevant information. The system then generates a Cypher query through a carefully designed prompt that includes both general Cypher syntax guidance and domain-specific instructions. The generated query is cleaned to remove any extraneous characters or formatting before execution against the knowledge graph.

To enhance the LLM’s understanding of the knowledge graph’s semantic structure, we enriched the schema with detailed comments for nodes, relationships, and properties. These schema comments provide context that helps the LLM interpret the meaning and purpose of each element within the graph. For nodes, we included descriptions explaining what each entity represents in the domain (e.g., “Protein: A biological macromolecule that performs various functions in organisms”). For relationships, we provided explanations of how entities interact (e.g., “hasFunction: Connects a protein to its biological function”). For properties, we described what information each attribute contains. These schema comments serve as semantic metadata that guides the LLM in selecting relevant pathways and constructing accurate queries, significantly reducing hallucinations when dealing with domain-specific terminology.

After completing the subtasks, the final Cypher query *C* is executed against the knowledge graph *G*. The results are then returned to the user in a readable format. Figure 2 shows the overall architecture of our proposed method.

**Figure 2.**
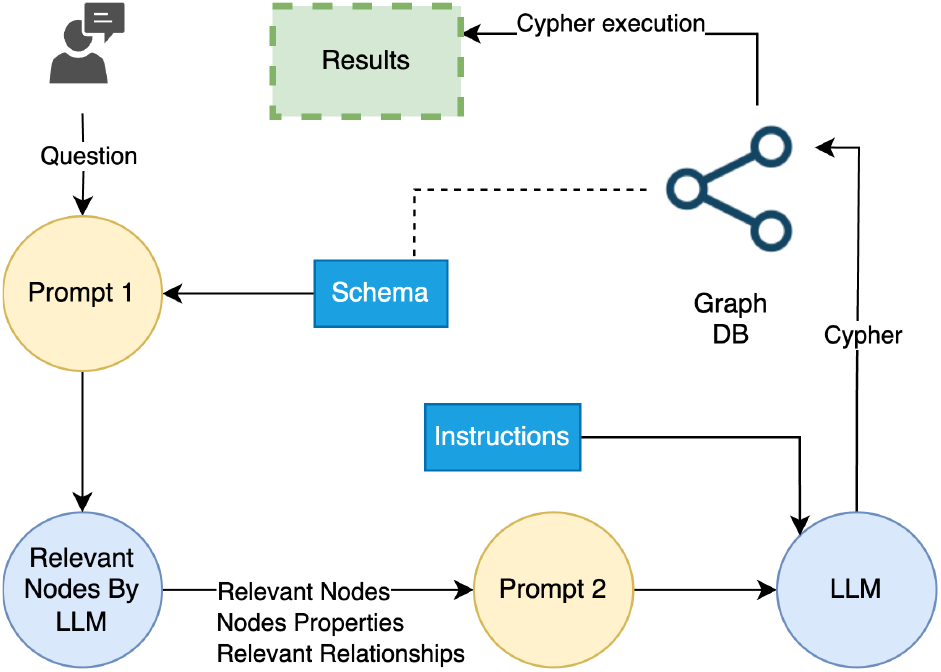
The architecture of our proposed approach. Prompt 1 instructs the system to find nodes relevant to the user query. Prompt 2 is consists of the list of those nodes, their properties, relevant relationships, and instructions to generate a Cypher query that answers the user’s question.

Prompt engineering is another powerful technique to guide LLMs toward more accurate outputs^44,45^. We developed both KG-agnostic and KG-specific instructions as part of our prompting strategy. For Cypher generation, we included general instructions such as: “Construct a valid Cypher query that answers the user’s question”, “Strictly follow the schema’s node labels, relationship types, and property names”, and “Use DISTINCT to avoid duplicate results whenever appropriate.” We also incorporated domain-specific instructions based on the target graph database. Critically, our approach integrates schema comments directly into the prompt, providing the LLM with essential semantic context about each node, relationship, and property. For example, when a user asks about “dark ion channels,” the prompt includes not just the node label “DarkIonChannel” but also its description: “Ion channels with limited experimental characterization.” This contextual enrichment helps the LLM understand domain-specific concepts that may not have been well-represented in its training data, substantially reducing hallucinations.

The full set of instructions used for prompt engineering is available in the supplementary data.

### Evaluation Metrics

To assess the effectiveness of our approach, we needed metrics that could meaningfully compare query results rather than just query syntax, since different Cypher queries can retrieve the same data through varied structures and variable names. We developed an evaluation framework based on set theory, treating query results as sets and computing standard information retrieval metrics that capture both the accuracy and completeness of the retrieved data.

Our evaluation framework centers on four complementary metrics, shown in Table 1. Precision measures the proportion of correctly retrieved results to the total retrieved results, indicating how accurate our generated queries are in returning only relevant information. While Precision focuses on accuracy, Recall measures completeness by calculating the proportion of correctly retrieved results to all relevant results that should have been retrieved. Since there is often a trade-off between Precision and Recall, we use the F1-score to provide a balanced measure of performance. As the harmonic mean of Precision and Recall, the F1-score penalizes extreme imbalances between these metrics. To complement these traditional metrics, we also employ the Jaccard Similarity coefficient, which provides a stringent measure of the overlap between the retrieved and relevant result sets. This metric is particularly useful for our context as it considers both the intersection and union of the result sets.

**Table 1.**
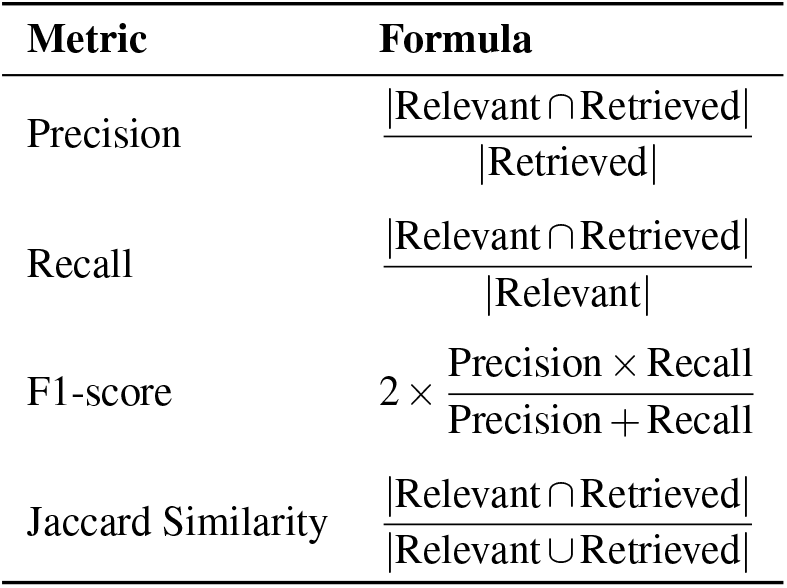
Evaluation Metrics Formulas.

This comprehensive evaluation framework allows us to assess both the correctness and completeness of our query results. The combination of these metrics provides a nuanced view of performance, where high scores across all metrics indicate that the generated queries not only retrieve the right information, but also avoid irrelevant results. In the context of knowledge graph querying, this is particularly important, as it ensures that users receive accurate and complete answers to their queries while minimizing noise in the results.

### User Validation

Given that LLMs can sometimes generate hallucinations, we implemented validation mechanisms to ensure the accuracy and relevance of generated Cypher queries. Our approach provides transparency by displaying both the generated Cypher query and a visual representation of its execution path. Since Cypher queries can be challenging for non-technical users to interpret, we developed a visualization technique that extracts execution plans from Neo4j and renders them as intuitive diagrams using NetworkX. This visualization highlights the nodes, relationships, and properties involved in the query, allowing users to visually verify whether the generated query aligns with their original intent. As shown in Figure 3, we visualize how P2C handles the query “List all the ions and gating mechanisms for the ‘K’ family of dark ion channels” by displaying the relevant nodes and relationships involved in the generated Cypher query. This visualization is created by parsing Neo4j’s execution plan^46^, extracting the query components, and rendering them graphically. This transparent approach builds user trust and provides an educational component that helps users understand how their natural language questions translate to graph database queries.

**Figure 3.**
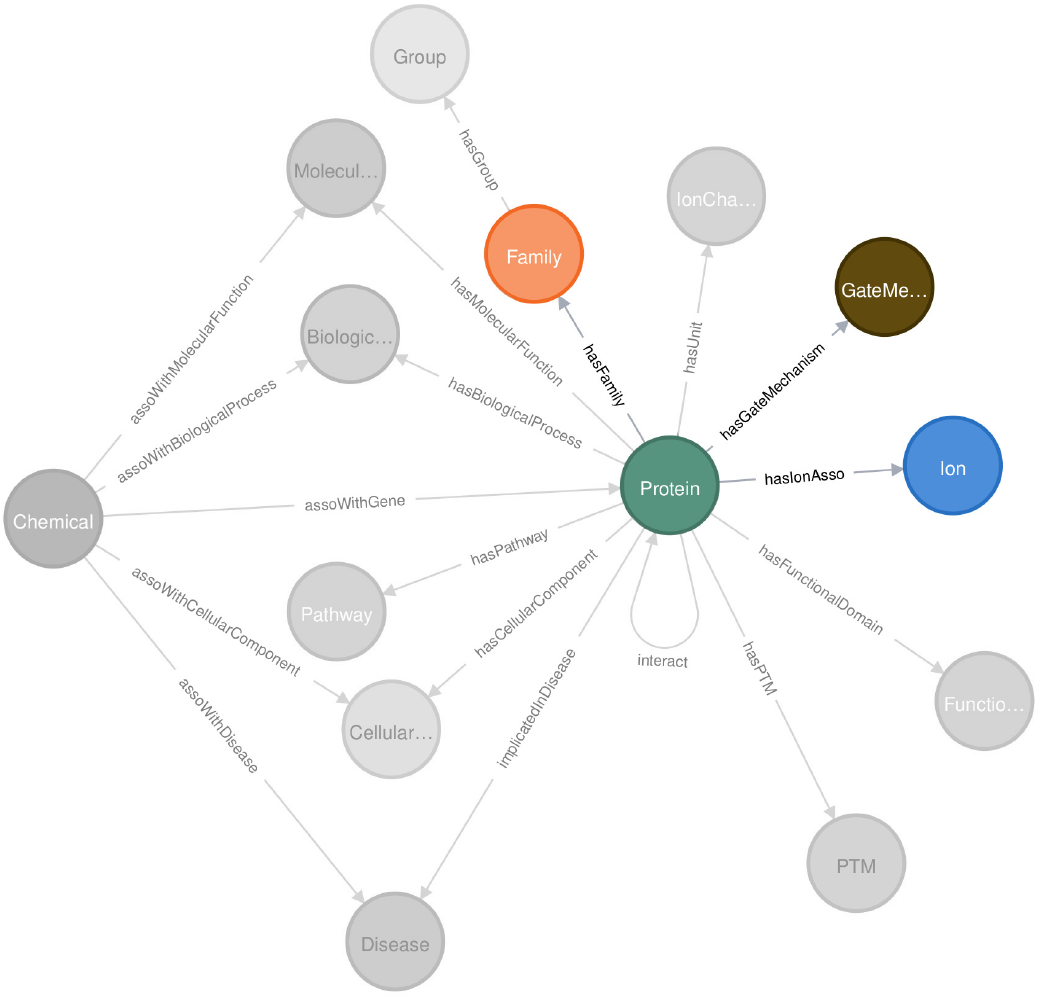
Displaying nodes and relationships from the schema that are involved in a generated Cypher query. This example visualizes the nodes P2C selects to answer “List all the ions and gating mechanisms for the ‘K’ family of dark ion channels”. The filtering process has reduced the schema complexity from 18 to 4 node types and from 18 to 3 relationship types, creating a tractable reasoning space for path discovery. The P2C generates and visualizes the following Cypher query by focusing on nodes and relationships involved: *“MATCH (dc:DarkIonChannel)-[:hasFamily]->(f:Family label: ‘K’) MATCH (dc)-[:hasIonAsso]->(i:Ion) MATCH (dc)-[:hasGateMechanism]->(g:GateMechanism) RETURN DISTINCT i*.*label AS Ion, g*.*label AS GatingMechanism LIMIT 100”*

## Results and Discussion

This section presents the results of our experiments evaluating the P2C approach for generating accurate Cypher queries from natural language inputs. We begin by generating Cypher queries using a baseline method for comparison, in which an LLM is directly prompted to generate Cypher queries using the crafted natural language test queries and the full schema of the KG. We then evaluate our proposed P2C approach in a similar manner using the same test queries. We compare the results obtained from executing the generated Cypher queries with those from the manually curated test queries, using the metrics described in the Methods section. Our findings demonstrate that P2C significantly improves query accuracy over the baseline method, validating the effectiveness of task splitting and prompt engineering in enhancing LLM performance for querying KGs.

### Baseline

As a starting point, we consider a baseline method where the LLM is prompted to generate a Cypher query directly from the user’s natural language query *Q*, the full schema of the knowledge graph *G*, and general instructions for Cypher query generation.

Figure 4 illustrates this baseline workflow using an example query written in natural language by a user. The user query is combined with Cypher generation instructions and the full schema of the ProKinO graph database, and provided to the LLM. However, due to the complexity and size of the schema, as well as potential ambiguities in the query, the LLM generates an incorrect Cypher query. Specifically, it uses a common property in KGs (i.e., “label”) instead of the correct property (i.e, “hasPrimaryName”) in this specific KG. Executing this incorrect query leads to no results, demonstrating the limitations of the baseline approach when dealing with large and complex KGs.

**Figure 4.**
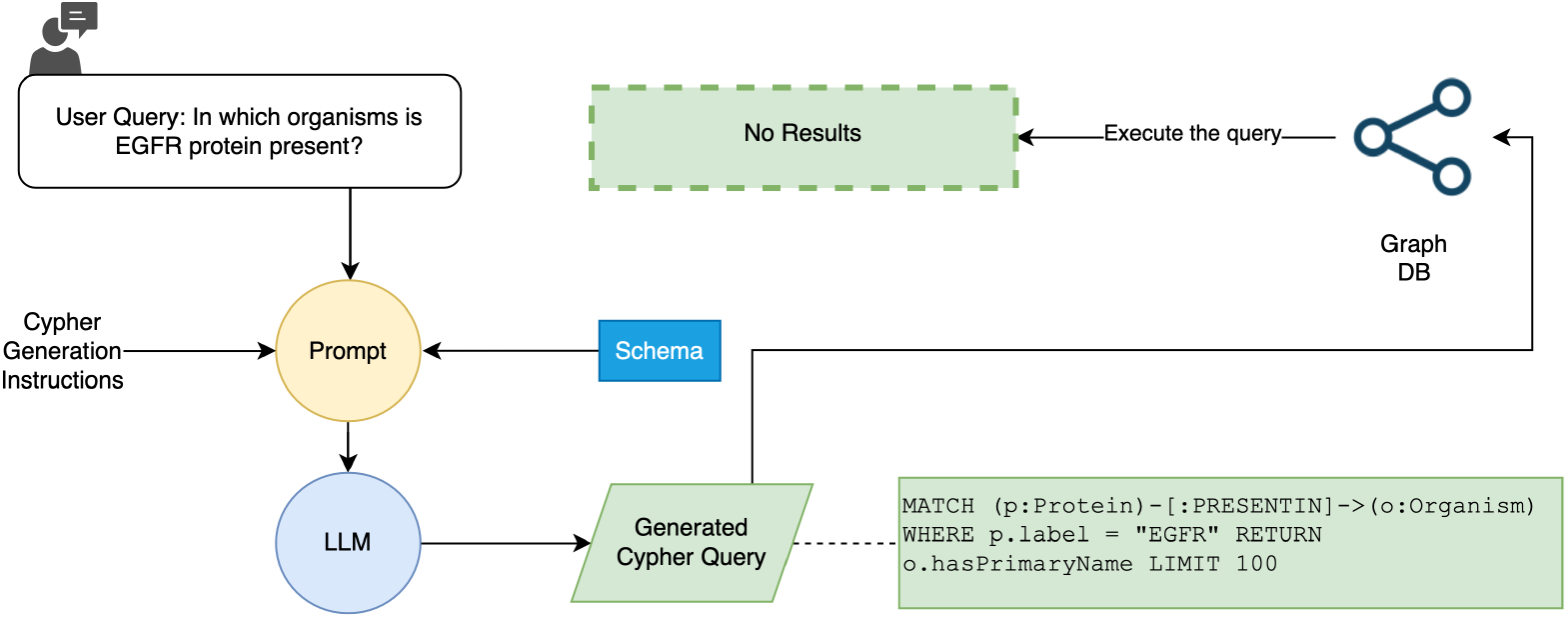
Baseline workflow for generating Cypher queries from natural language inputs. The user query “In which organisms is EGFR protein present?”, is combined with general instructions and the schema of the ProKinO graph database and provided to the LLM. The LLM generates a semantically incorrect Cypher query resulting in no data retrieved upon execution. It demonstrates the challenges of generating queries with LLMs when the context size is large and contains irrelevant extra information.

This highlights the baseline’s limitations in handling large and complex KGs, necessitating a more effective method. These limitations prompted the development of our Prompt2Cypher (P2C) approach, which we introduce in the following subsection.

### Prompt2Cypher Improves Performance

Prompt2Cypher (P2C) addresses the limitations of the baseline method by introducing task splitting and prompt engineering. Task splitting breaks down the complex process of generating a Cypher query into manageable subtasks, allowing the LLM to focus on specific aspects of the query.

Prompt engineering involves crafting detailed instructions and providing relevant context to guide the LLM’s output, ensuring alignment with the KG’s schema. Figure 5 illustrates the P2C workflow. In the first step, the user query is processed to identify relevant nodes based on the schema. Subsequent steps involve extracting properties and relationships pertinent to these nodes, which are then used to construct accurate Cypher queries.

**Figure 5.**
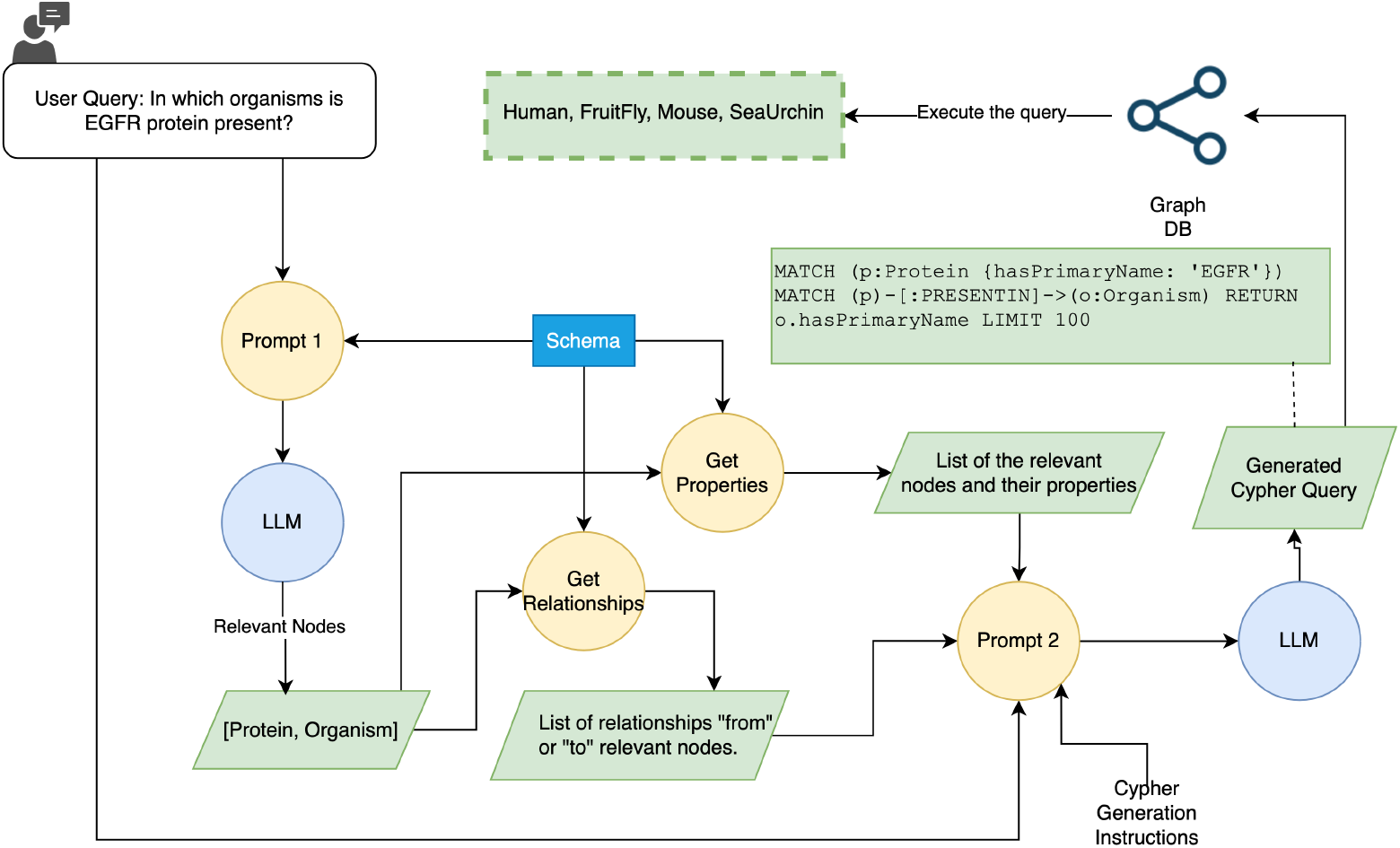
Proposed P2C workflow for generating Cypher queries (based on Figure 2). The process involves two prompts. In Prompt 1, the user query is processed to identify relevant nodes based on the schema of the graph database. In this example, the relevant nodes are Protein and Organism. Then, in Get Properties and Get Relationship functions the schema is filtered out to contain only properties and relationships related to these nodes. This information is used to construct Prompt 2. Prompt 2 integrates the properties and relationships to form a more accurate Cypher query. The generated query correctly uses the property “hasPrimaryName” for the Protein entity. When executed against the graph database, this query successfully returns the list of organisms (Human, FruitFly, Mouse, SeaUrchin) containing the EGFR protein.

By combining these strategies, P2C aims to enhance query accuracy and reduce the occurrence of hallucinations, making complex KGs more accessible to non-technical users. To assess the effectiveness of P2C, we conducted comprehensive evaluations described in the next subsection.

### Evaluations

To evaluate the effectiveness of P2C, we conducted experiments using two domain-specific KGs: ProKinO and the ICKG. We formulated a set of 30 test queries for each KG, varying in complexity from 1-hop to 3-hop queries. These queries were designed to assess the system’s ability to handle different levels of query complexity. We compared the performance of P2C with the baseline using standard evaluation metrics of precision, recall, F1 score, and Jaccard similarity. For both KGs, we tested our approach and the baseline with 30 test queries by three different OpenAI’s models: GPT-4o-mini, o1-mini, and o1-preview.

We utilized the workflows illustrated in Figures 4 and 5 to evaluate both the baseline and our proposed approach, P2C. Alongside the user query and schema, both the baseline and P2C workflows incorporate a prompt of Cypher generation instructions, which guide the LLM in crafting a well-structured Cypher query to respond to the user query. For consistency, a limit of 100 results was set for evaluation, as some queries could return thousands of entries, complicating comparisons between baseline results and those generated by P2C.

The comparative analysis reveals distinct performance patterns across LLMs and between the two knowledge graphs, as shown in Table 2. Among the tested models, o1-preview demonstrated superior performance, achieving the highest scores across all metrics for both knowledge graphs. For ICKG, o1-preview with P2C achieved a precision of 0.9467 and recall of 0.8832, significantly outperforming both gpt-4o-mini (precision: 0.8532, recall: 0.7940) and o1-mini (precision: 0.8755, recall: 0.8196). Similarly for ProKinO, o1-preview with P2C showed remarkable performance with precision of 0.8939 and recall of 0.9483, compared to gpt-4o-mini (precision: 0.7350, recall: 0.6791) and o1-mini (precision: 0.8333, recall: 0.8292).

**Table 2.**
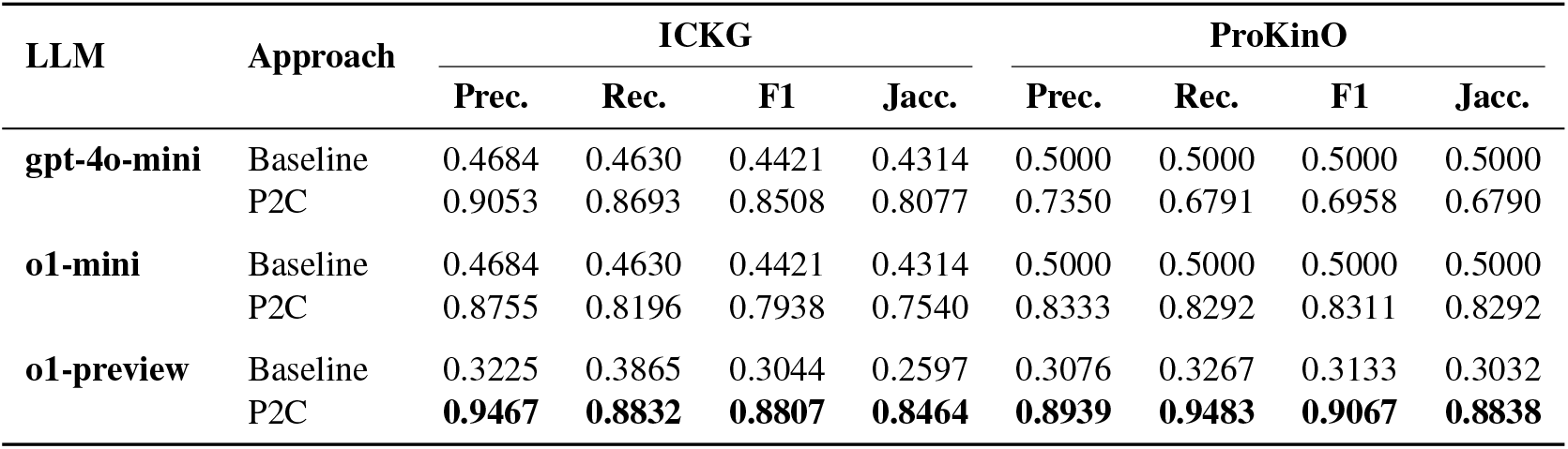
Performance Metrics for ICKG and ProKinO Test Queries.

| LLM | Approach | ICKG |  |  |  | ProKinO |  |  |  |
| --- | --- | --- | --- | --- | --- | --- | --- | --- | --- |
|  |  | Prec. | Rec. | F1 | Jacc. | Prec. | Rec. | F1 | Jacc. |
| **gpt-4o-mini** | Baseline | 0.4684 | 0.4630 | 0.4421 | 0.4314 | 0.5000 | 0.5000 | 0.5000 | 0.5000 |
|  | P2C | 0.9053 | 0.8693 | 0.8508 | 0.8077 | 0.7350 | 0.6791 | 0.6958 | 0.6790 |
| **o1-mini** | Baseline | 0.4684 | 0.4630 | 0.4421 | 0.4314 | 0.5000 | 0.5000 | 0.5000 | 0.5000 |
|  | P2C | 0.8755 | 0.8196 | 0.7938 | 0.7540 | 0.8333 | 0.8292 | 0.8311 | 0.8292 |
| **o1-preview** | Baseline | 0.3225 | 0.3865 | 0.3044 | 0.2597 | 0.3076 | 0.3267 | 0.3133 | 0.3032 |
|  | P2C | **0.9467** | **0.8832** | **0.8807** | **0.8464** | **0.8939** | **0.9483** | **0.9067** | **0.8838** |

The performance difference between ICKG and ProKinO is particularly noteworthy. For the o1-preview with P2C, while ICKG showed higher precision (0.9467 vs 0.8939), ProKinO demonstrated better recall (0.9483 vs 0.8832). This variation can be attributed to several factors. First, the structural differences between the two knowledge graphs play a key role; ICKG has a more focused domain with 18 node types and relationship types, while ProKinO has a more complex structure with 47 node types and 37 relationship types. The focused nature of ICKG likely contributes to higher precision in query generation, as the model has fewer options to consider when constructing queries. ProKinO’s higher recall suggests that despite its complexity, its comprehensive schema actually helps the model capture more relevant results. This could be because ProKinO’s richer network of relationships provides multiple paths to reach relevant information, increasing the likelihood of retrieving all pertinent results. Additionally, since test queries were created manually, variations in query complexity between the two KGs could have influenced these metrics.

While our method consistently outperforms the baseline across most metrics, we observed occasional decreases in Recall, F1, and Jaccard scores. This discrepancy arises not from incorrect data retrieval, but rather from differences in the properties returned by the generated queries compared to the ground truth. In some cases, the ground truth queries explicitly specify returning both entities and their associated attributes, whereas the generated queries may only return the entities themselves. For example, a ground truth query might return both a protein and the biological process it participates in, whereas the generated query might return only the protein name as shown in Figure 4. Similarly, another ground truth query may list both an ion channel and the pathway it belongs to, while the generated query only reports the ion channel label, because there were no explicit request on what to return in the question. These subtle variations in output format, rather than errors in the retrieved results, lead to lower scores in certain evaluation metrics despite the generated queries retrieving correct and relevant information.

To further understand performance variations, we analyzed how P2C’s effectiveness changes with query complexity. We categorized test queries into three complexity levels: 1-hop (simple), 2-hop (medium), and 3-hop (complex) queries, based on the number of node traversals required. Table 3 presents performance metrics across these categories for both knowledge graphs.

**Table 3.**
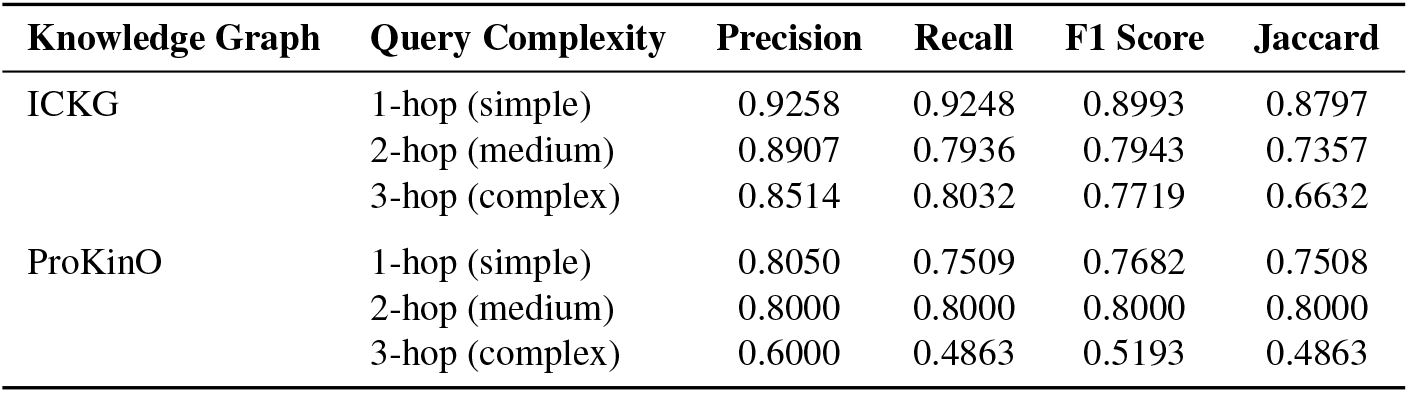
Performance Metrics by Query Complexity.

Figure 6 visualizes these performance trends across query complexity levels. For ICKG, we observe a gradual decline in performance as complexity increases, with F1 scores decreasing from 0.8993 for 1-hop queries to 0.7719 for 3-hop queries. Despite this decline, precision remains above 0.85 even for complex queries, indicating that P2C maintains relatively high accuracy despite increased query complexity.

**Figure 6.**
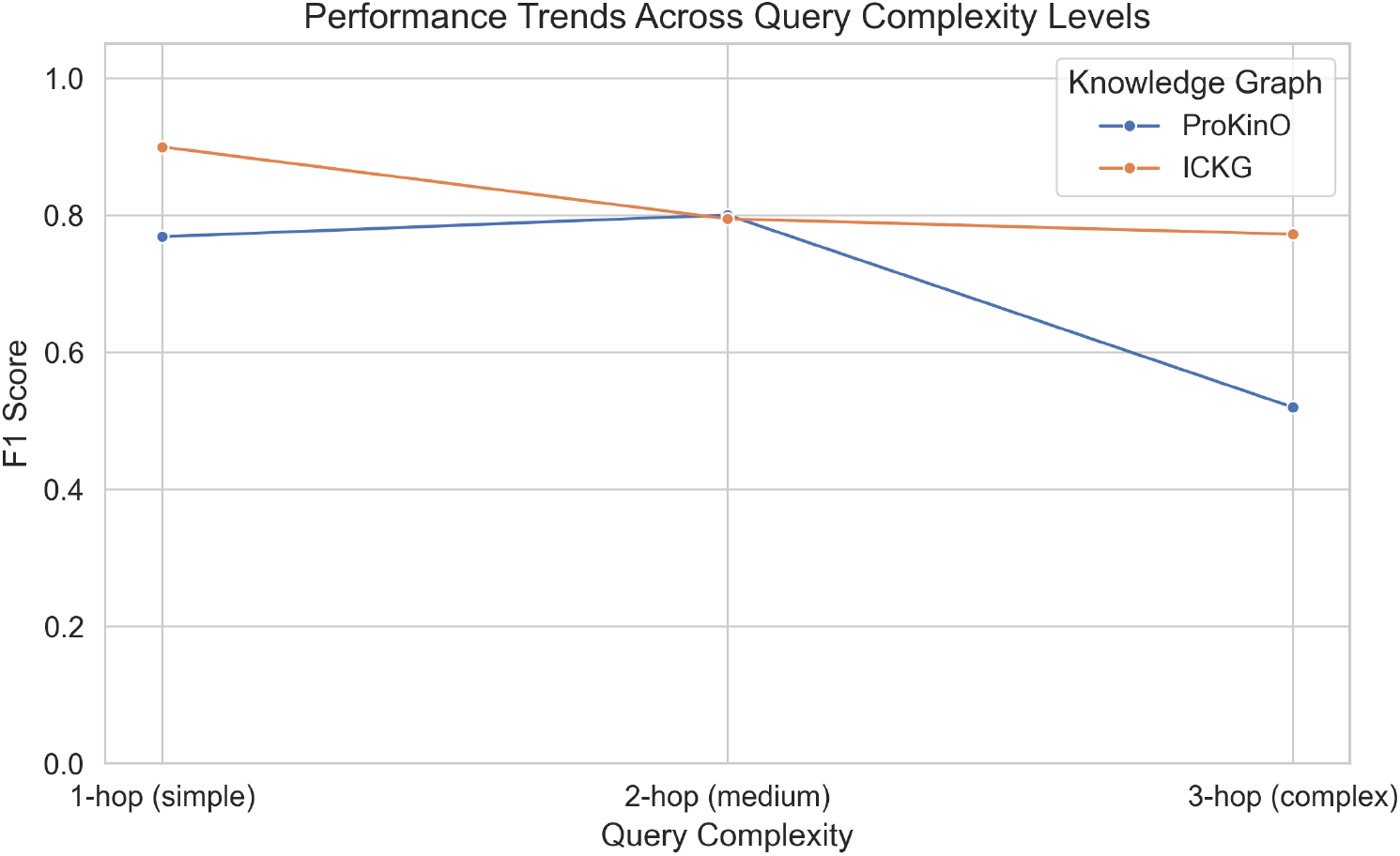
Performance trends of P2C across query complexity levels for both knowledge graphs. The figure illustrates how performance degrades more rapidly for the more complex ProKinO schema compared to the more focused ICKG schema as query complexity increases.

For ProKinO, the impact of complexity is more pronounced, with F1 scores showing stability between simple and medium complexity queries (0.7682 and 0.8000, respectively), but dropping significantly to 0.5193 for complex queries. This steeper decline for ProKinO compared to ICKG can be attributed to its greater schema complexity (47 node types and 37 relationship types versus ICKG’s 18 node and relationship types), which amplifies the challenge of multi-hop reasoning. These findings suggest that while P2C handles simple and medium complexity queries effectively across different knowledge graph schemas, its performance for complex multi-hop queries is more schema-dependent.

### Ablation Study

To better understand the contribution of each component in the Prompt2Cypher framework, we conducted a comprehensive ablation study across both knowledge graphs. This analysis evaluates the impact of three key components: (1) specialized instructions in prompts, (2) schema comments that provide context about node and relationship types, and (3) relevant node filtering that narrows the search space. For each component, we measured performance when the component was removed from the full system. These findings complement our complexity analysis (Table 3), which showed performance degradation with increasing query complexity, particularly for ProKinO.

Table 4 shows the results of our ablation study using GPT-4o-mini on both the ProKinO and ICKG datasets. We present the actual performance metrics of each configuration, allowing direct comparison between the complete system and versions with individual components removed.

**Table 4.** Ablation Study Results for ICKG and ProKinO.

| Ablation | ICKG |  |  | ProKinO |  |  |
| --- | --- | --- | --- | --- | --- | --- |
|  | Prec. | Rec. | F1 | Prec. | Rec. | F1 |
| Complete System | 0.905 | 0.869 | 0.850 | 0.735 | 0.679 | 0.695 |
| Without Instructions | 0.662 | 0.650 | 0.655 | 0.410 | 0.467 | 0.427 |
| Without Schema Comments | 0.443 | 0.433 | 0.435 | 0.467 | 0.450 | 0.456 |
| Without Relevant Nodes | 0.235 | 0.194 | 0.211 | 0.767 | 0.767 | 0.767 |

As shown in Table 4, the impact of removing components varies significantly between the two knowledge graphs. For ProKinO, removing specialized instructions results in a significant performance drop (F1 score dropping from 0.695 to 0.427), while for ICKG, removing relevant node filtering has the most dramatic impact (F1 score dropping from 0.850 to 0.211). Interestingly, removing the relevant nodes filtering actually improves overall performance for ProKinO (F1 score of 0.767), suggesting that for this complex schema, considering all nodes sometimes provides better coverage than selective filtering. In contrast, for ICKG, removing relevant node filtering has the most dramatic negative impact (F1 score dropping from 0.850 to 0.211), indicating that for more focused schemas, narrowing the search space is critical. These contrasting results highlight how knowledge graph complexity influences the importance of different components.

The differential impact of component removal reveals how knowledge graph complexity influences the importance of each component in P2C: more complex schemas like ProKinO rely heavily on instructions and schema context, while more focused schemas like ICKG benefit most from narrowing the search space through relevant node filtering. These findings demonstrate that the effectiveness of P2C stems from the synergistic integration of all three components, with their relative importance varying based on knowledge graph complexity. The specialized instructions guide the LLM toward valid Cypher syntax, schema comments provide essential context about the knowledge graph structure, and relevant node filtering helps narrow the search space.

The contrasting ablation results between ProKinO and ICKG suggest that the optimal prompting strategy should be adapted to the complexity of the underlying knowledge graph. For larger, more complex schemas like ProKinO, comprehensive instructions and schema context are paramount, while for more focused schemas like ICKG, the ability to identify relevant nodes becomes the limiting factor. This suggests that future prompt engineering approaches might benefit from dynamic adjustment based on knowledge graph complexity, potentially implementing more sophisticated node filtering approaches for complex knowledge graphs like ProKinO. This ablation study highlights the adaptability of our approach to different knowledge graph structures, suggesting that future improvements to the framework could potentially employ adaptive approaches that emphasize different components based on knowledge graph characteristics.

## Conclusion

In this study, we presented Prompt2Cypher (P2C), a novel approach that uses LLMs to generate accurate Cypher queries from natural language inputs for domain-specific knowledge graphs. By employing task splitting and prompt engineering, P2C effectively addresses the challenges of querying complex KGs, as demonstrated by its superior performance compared to a baseline method across various evaluation metrics. These results highlight the potential of LLMs in making structured data more accessible to both technical and non-technical users. The success of P2C can be attributed to the combination of task decomposition, which simplifies the query generation process into manageable subtasks, and comprehensive prompt engineering, which provides detailed instructions that align the LLM’s outputs with the graph schema’s constraints. This strategy enhances the performance of the LLM and makes the querying process more accessible to non-technical users, bridging the gap between advanced data structures and user-friendly interfaces.

Our ablation study revealed important insights about how different components of P2C contribute to its effectiveness across knowledge graphs of varying complexity. For complex schemas like ProKinO, specialized instructions and schema comments proved to be the most critical components, demonstrating that semantic context is essential for navigating large, intricate knowledge structures. In contrast, for more focused schemas like ICKG, relevant node filtering emerged as the dominant factor, highlighting the importance of narrowing the search space in domains with well-defined entity relationships. Furthermore, we observed that the relative importance of these components shifts with query complexity—for simple queries, schema comments provide sufficient guidance, while complex multi-hop queries benefit most from relevant node filtering to manage the exponentially growing search space. These findings suggest that an adaptive approach to prompt engineering, which dynamically emphasizes different components based on both schema complexity and query type, could further enhance LLM performance in knowledge graph querying.

Our analysis of query complexity further revealed that P2C’s performance is more resilient to increasing query complexity in knowledge graphs with focused schemas (like ICKG) compared to those with complex schemas (like ProKinO). As queries require traversing multiple relationships (3-hop queries), the compounding complexity of larger schemas becomes a significant challenge, with F1 scores dropping more dramatically for ProKinO (from 0.7682 to 0.5193) than for ICKG (from 0.8993 to 0.7719).

The dramatic performance gap between general-purpose approaches like Text2Cypher^25^ and our method underscores the unique challenges posed by domain-specific knowledge graphs. While Text2Cypher has shown success with simpler schemas, its complete failure (0% accuracy) on our biomedical knowledge graphs highlights the necessity of specialized approaches for complex scientific domains. This finding suggests that future work in natural language query generation must carefully consider domain-specific requirements and schema complexity when developing new approaches.

By enabling users to ask questions in their own language, P2C could be used to develop chatbots that can answer questions about domain-specific knowledge and provide personalized recommendations. P2C empowers researchers and scientists to discover new insights, formulate new hypotheses, and make informed decisions based on the wealth of information stored in KGs. This capability can be particularly valuable in domains such as biology and medicine, where complex KGs are used to represent vast amounts of data related to genes, proteins, diseases, and pathways.

Despite its promise, P2C faces challenges with multi-hop queries and highly complex or large KGs. This is particularly evident in the performance drop for complex multi-hop queries in rich schemas like ProKinO, as shown in our complexity analysis. For example, we encountered difficulties integrating the Protein Ontology (PRO) with P2C due to its OWL-based structure, which is not readily amenable to intuitive Cypher querying. To address this, future work could explore preprocessing steps to restructure such ontologies into more LPG (labeled property graph)-friendly schemas. This would involve extracting named entities, simplifying relationships, and creating explicit node labels and direct connections to facilitate more intuitive query generation by LLMs.

Furthermore, the zero-shot nature of our experiments suggests that few-shot learning or more targeted prompting could further improve performance. Future research could also investigate incorporating retrieval-augmented generation or fine-tuning strategies to enhance P2C’s capabilities and address limitations in handling complex queries. Exploring pathfinding algorithms and incorporating user feedback mechanisms could enhance the system’s adaptability and robustness.

Despite these challenges, P2C represents a significant step forward in facilitating knowledge discovery and data mining by bridging the gap between complex data structures and user-friendly interfaces. This approach holds promise for advancing research and applications that rely on rich, structured data representations in various domains.

## Acknowledgements (not compulsory)

Funding for NK from the Illuminating the Druggable Genome NIH Common Fund is acknowledged (U01 CA239106/CA/NCI NIH; U01 CA271376/CA/NCI).

## Author contributions statement

S.S. conceived and designed the methodology, implemented the P2C system, conducted experiments, and performed data analysis. N.G. contributed to data collection and validation, particularly for the ion channels knowledge graph. K.K. provided expertise in knowledge graphs, validated evaluation metrics, and supervised the methodological aspects including graph query generation and results analysis. N.K. provided domain expertise, supervised the overall project direction, validated the biological aspects of the ion channels knowledge graph, and ensured quality control of the research. All authors contributed to writing and reviewing the manuscript.

## Data Availability

The datasets generated and analyzed during the current study, along with the source code and supplementary materials, are available in the Prompt2Cypher repository: https://github.com/noghte/prompt2cypher. All data and materials are publicly accessible under an open-source license to ensure reproducibility and further research.

## Supplementary Information

The following supplementary data for both knowledge graphs ProKinO and ICKG are available:

1. **Supplementary Data S1**: Cypher generation instructions
2. **Supplementary Data S2**: Schema files
3. **Supplementary Data S3**: Complete list of test queries used for evaluating P2C and baseline approaches
4. **Supplementary Data S4**: Raw evaluation results for all test queries
5. **Supplementary Data S5**: Source code and implementation details

All supplementary data and code are available at: https://github.com/noghte/prompt2cypher

## References

1. Zhang, Y. et al. Biokg: a comprehensive, large-scale biomedical knowledge graph for ai-powered, data-driven biomedical research. bioRxiv (2023).

2. Abu-Salih, B. Domain-specific knowledge graphs: A survey. J. Netw. Comput. Appl. 185, 103076 (2021).

3. Soleymani, S. et al. Dark kinase annotation, mining, and visualization using the protein kinase ontology. PeerJ 11, e16087 (2023).

4. Taujale, R. et al. Informatic challenges and advances in illuminating the druggable proteome. Drug discovery today 29, 103894 (2024).

5. Androutsopoulos, I., Ritchie, G. D. & Thanisch, P. Natural language interfaces to databases–an introduction. Nat. language engineering 1, 29–81 (1995).

6. Templeton, M. & Burger, J. F. Problems in natural-language interface to dbms with examples from eufid. In First Conference on Applied Natural Language Processing, 3–16 (1983).

7. Popescu, A.-M., Etzioni, O. & Kautz, H. Towards a theory of natural language interfaces to databases. In Proceedings of the 8th international conference on Intelligent user interfaces, 149–157 (2003).

8. Kargar, M., Zihayat, M. & Szlichta, J. Mining and exploration of attributed graphs: theory and applications. In Proceedings of the 29th Annual International Conference on Computer Science and Software Engineering, 397–398 (2019).

9. Fan, Y. et al. Metasql: A generate-then-rank framework for natural language to sql translation. In 2024 IEEE 40th International Conference on Data Engineering (ICDE), 1765–1778 (IEEE, 2024).

10. Zettlemoyer, L. S. & Collins, M. Learning to map sentences to logical form: Structured classification with probabilistic categorial grammars. arXiv preprint arXiv:1207.1420 (2012).

11. Liu, M. & Xu, J. Nli4db: A systematic review of natural language interfaces for databases. arXiv preprint arXiv:2503.02435 (2025).

12. Li, F. & Jagadish, H. V. Constructing an interactive natural language interface for relational databases. Proc. VLDB Endow. 8, 73–84 (2014).

13. Dubey, A. et al. The llama 3 herd of models. arXiv preprint arXiv:2407.21783 (2024).

14. Fakhoury, S., Naik, A., Sakkas, G., Chakraborty, S. & Lahiri, S. K. Llm-based test-driven interactive code generation: User study and empirical evaluation. arXiv preprint arXiv:2404.10100 (2024).

15. Hayes, T. et al. Simulating 500 million years of evolution with a language model. bioRxiv 2024–07 (2024).

16. Yao, J.-Y., Ning, K.-P., Liu, Z.-H., Ning, M.-N. & Yuan, L. Llm lies: Hallucinations are not bugs, but features as adversarial examples. arXiv preprint arXiv:2310.01469 (2023).

17. Tonmoy, S. et al. A comprehensive survey of hallucination mitigation techniques in large language models. arXiv preprint arXiv:2401.01313 (2024).

18. Martino, A., Iannelli, M. & Truong, C. Knowledge injection to counter large language model (llm) hallucination. In European Semantic Web Conference, 182–185 (Springer, 2023).

19. Beheshti, M. et al. Evaluating the reliability of chatgpt for health-related questions: A systematic review. Informatics 12, DOI: 10.3390/informatics12010009 (2025).

20. Soman, K. et al. Biomedical knowledge graph-optimized prompt generation for large language models. Bioinformatics 40, btae560 (2024).

21. Xu, X., Liu, C. & Song, D. Sqlnet: Generating structured queries from natural language without reinforcement learning. arXiv preprint arXiv:1711.04436 (2017).

22. Zhong, V., Xiong, C. & Socher, R. Seq2sql: Generating structured queries from natural language using reinforcement learning. CoRR abs/1709.00103 (2017).

23. Wang, B., Shin, R., Liu, X., Polozov, O. & Richardson, M. Rat-sql: Relation-aware schema encoding and linking for text-to-sql parsers. arXiv preprint arXiv:1911.04942 (2019).

24. Guo, A., Li, X., Xiao, G., Tan, Z. & Zhao, X. Spcql: A semantic parsing dataset for converting natural language into cypher. In Proceedings of the 31st ACM International Conference on Information & Knowledge Management, 3973–3977 (2022).

25. Ozsoy, M. G., Messallem, L., Besga, J. & Minneci, G. Text2cypher: Bridging natural language and graph databases. arXiv preprint arXiv:2412.10064 (2024).

26. Ozsoy, M. G. Neo4j text2cypher: Analyzing model struggles and dataset improvements. https://medium.com/neo4j/neo4j-text2cypher-analyzing-model-struggles-and-dataset-improvements-0b965fd3ebfa (2025). Accessed: 2025-04-19.

27. Sivasubramaniam, S., Osei-Akoto, C. E., Zhang, Y., Stockinger, K. & Fuerst, J. Sm3-text-to-query: Synthetic multi-model medical text-to-query benchmark. Adv. Neural Inf. Process. Syst. 37, 88627–88663 (2024).

28. Zhong, Z. et al. Synthet2c: Generating synthetic data for fine-tuning large language models on the text2cypher task. arXiv preprint arXiv:2406.10710 (2024).

29. Shekarpour, S. et al. Generating sparql queries using templates. Web Intell. Agent Syst. An Int. J. 11, 283–295 (2013).

30. Baraki, W. W. Leveraging large language models for accurate cypher query generation: Natural language query to cypher statements (2024).

31. Zhang, Y. et al. Making large language models perform better in knowledge graph completion. arXiv preprint arXiv:2310.06671 (2023).

32. Shu, D. et al. Knowledge graph large language model (kg-llm) for link prediction. arXiv preprint arXiv:2403.07311 (2024).

33. Alqaaidi, S. K., Bozorgi, E. & Kochut, K. J. Multiple relations classification using imbalanced predictions adaptation. arXiv preprint arXiv:2309.13718 (2023).

34. Jiang, J. et al. Kg-agent: An efficient autonomous agent framework for complex reasoning over knowledge graph. arXiv preprint arXiv:2402.11163 (2024).

35. Choi, S. R. & Lee, M. Transformer architecture and attention mechanisms in genome data analysis: a comprehensive review. Biology 12, 1033 (2023).

36. Marvin, G., Hellen, N., Jjingo, D. & Nakatumba-Nabende, J. Prompt engineering in large language models. In International conference on data intelligence and cognitive informatics, 387–402 (Springer, 2023).

37. Sheils, T. K. et al. Tcrd and pharos 2021: mining the human proteome for disease biology. Nucleic Acids Res. 49, D1334–D1346 (2021).

38. Uniprot: the universal protein knowledgebase in 2025. Nucleic Acids Res. 53, D609–D617 (2025).

39. Szklarczyk, D. et al. The string database in 2023: protein–protein association networks and functional enrichment analyses for any sequenced genome of interest. Nucleic acids research 51, D638–D646 (2023).

40. Aleksander, S. A. et al. The gene ontology knowledgebase in 2023. Genetics 224, iyad031 (2023).

41. Milacic, M. et al. The reactome pathway knowledgebase 2024. Nucleic acids research 52, D672–D678 (2024).

42. Carbon, S. & Mungall, C. Gene ontology data archive, DOI: 10.5281/zenodo.14083199 (2024). Dataset.

43. Xiong, C., Liu, C., Li, H. & Li, X. Hlspilot: Llm-based high-level synthesis. arXiv preprint arXiv:2408.06810 (2024).

44. Polak, M. P. & Morgan, D. Extracting accurate materials data from research papers with conversational language models and prompt engineering. Nat. Commun. 15, 1569 (2024).

45. Maharjan, J. et al. Openmedlm: prompt engineering can out-perform fine-tuning in medical question-answering with open-source large language models. Sci. Reports 14, 14156 (2024).

46. Neo4j. Understanding Execution Plans (2025). Accessed: 2025-01-23.

